# Inferring physical cell-cell communication networks from scRNAseq data using univariate linear models

**DOI:** 10.1101/2025.07.07.663470

**Authors:** Sodiq Ayobami Hameed, Luis Fernando Iglesias-Martinez, Walter kolch, Vadim Zhernovkov

## Abstract

Cells in tissues interact by direct physical contact or over short and long distances via secreted mediators. Cell-cell communication inference has now become routine in downstream scRNAseq analysis but this mostly fails to capture physical cell-cell interactions due to tissue dissociation. Multiplets (mostly doublets) in scRNAseq may represent undissociated physically attached cells that become sequenced together. Hence, identifying multiplets may serve as a good starting point to harness scRNAseq data for physical cell-cell interaction inference. In this study, we develop a computational method which utilizes univariate linear models (ULM) to identify multiplets in scRNAseq datasets, predict their cellular compositions, and infer physical cell-cell interaction networks. Indeed, our method showed good sensitivity (∼56%) with excellent precision (∼99%) in predicting FACS sorted doublet cell pairs with known constituents (ground truth), recording comparable or superior performance over two other existing methods. Also, applying it to scRNAseq data of partially dissociated tissues containing real multiplets unraveled physical networks which recapitulated the microanatomical structures of the tested tissues. This further underscores the accuracy in our predictions to capture biologically meaningful interactions. Finally, we tested our method on classical scRNAseq datasets and obtained biologically reasonable results. For example, when tested on classical cancer scRNAseq datasets, we recovered important interactions which followed biologically plausible cell interactions, validated by cell-cell colocalization in matched spatial transcriptomics datasets. This reassured the accuracy of our method in depicting physical interactions only between cells that were truly in close proximity in tissues.

## 1.0 INTRODUCTION

Cells in body tissues exist in a complex and dynamic multicellular environment with neighboring cells where each cell’s function is influenced by the network of interactions with other cells [1]. Cells in multicellular organisms continuously communicate from early embryonic development to adult life through these dynamic interaction networks. Whenever these interactions are perturbed, cellular functions may break down, leading to diseases and multiple pathogenic scenarios. For example, tumor development and progression involve complex interactions and dynamic co-evolution between neoplastic cells, immune cells and stromal cells [2]. Indeed, the spatial context of cellular organization and the associated cell-cell communications in tissues determines physiological processes ranging from cell differentiation, proliferation, and immune response to homeostasis [3].

Single-cell RNA sequencing (scRNAseq) is a powerful technique which allows massive parallel profiling of thousands of cells to unravel cellular function and heterogeneity in health and disease [4]. Cell-cell communication inference has now become a routine method in scRNAseq data processing. Cells are clustered together based on markers, and the expression of matching ligands and receptors are assessed to infer interaction events [5]. Multiple tools exist for this purpose [6–10]. However, since scRNAseq protocols involve complete tissue dissociation and homogenization before sequencing, the spatial context of cell-cell interactions is disrupted. This limits the aforementioned scRNAseq tools to map cell neighborhoods in the native context within tissues [11]. This issue is surmounted by spatial transcriptomics techniques which allow gene expression in cells to be interrogated in their spatial tissue context, thereby improving the accuracy of cell interaction inference [12]. However, traditional spatial transcriptomics cannot localize RNA transcripts precisely at single-cell resolution, rather they capture a region of 10-100 cells in a spatial location in tissue and aggregate the genes originating from these cells [13]. In addition, newer techniques which perform spatial transcriptomics at true single-cell resolution can only localize a few hundreds to thousands of preselected genes per cell [14–17], thereby limiting unbiased applications. There are concerted efforts towards the utilization of modified scRNAseq techniques where tissues are partially dissociated to generate intentional cell aggregates prior to sequencing for native physical interaction inference [1,18–20]. These methods are efficient in deciphering physical cell interactions, however, they are technically demanding and require specialized personnel and equipment [21].

Multiplets in conventional droplet-based scRNAseq represent cell aggregates, which occur when more than one cell is captured in a single reaction droplet and sequenced together in a single barcode. These result largely from undissociated cells which remain physically attached to each other during library preparation or occasionally from the random co-encapsulation of cells [3]. Randomly generated multiplets can be largely avoided by careful dilution of cell suspensions during library preparation. However, the undissociated fractions usually represent biologically interactings cells which cannot be eliminated by sample dilution [22]. Hence, identifying multiplets may serve as a good starting point to harness scRNAseq data for physical cell-cell interaction inference. In this study, we utilized univariate linear models to identify multiplets in scRNAseq datasets and predict their cellular composition to infer physical cell-cell interaction networks in tissues. We tested this method on doublet cell pairs, partially dissociated tissues and conventional scRNAseq datasets of healthy and cancer tissues. We also validated the identified networks from scRNAseq using matched spatial transcriptomics dataset and downstream ligand-receptor analyses.

## 2.0 METHODS

### 2.1 Single-cell RNA sequencing and spatial transcriptomics data analysis

Preprocessed publicly available scRNAseq datasets were obtained, some of which were completely dissociated (singlet dataset) and some partially dissociated (doublets/ multiplets). Where applicable, the associated spatial transcriptomics datasets were also obtained. Data were processed using the Seurat package v5.1.1 in R v4.2.2. The scRNAseq data were preprocessed to include only cells containing a minimum of 500 genes, less than 25 percent mitochondrial genes, and less than 5000 total UMI. The count matrix was log2 transformed and the top 2,000 highly variable genes (HVG) were used for downstream analysis. A principal component analysis (PCA) was performed on the top 2,000 HVGs, and the top 20-30 principal components (PCs) were selected based on visualization from an elbow plot of the first 50 PCs to generate a uniform manifold approximation and projection (UMAP) plot. Then, cells were clustered using the nearest neighbor clustering algorithm at a resolution of 0.3-0.35.

#### 2.1.1 Dendritic cell and T cell singlet and doublet scRNAseq datasets

The scRNAseq count matrix and the associated metadata including cell labels consisting of single-positive dendritic cells (DC), single-positive T cells and double positive DC-T pairs from Giladi et al. [23] were obtained from the GEO database (GSE135382). Cells were assigned cell type labels by transferring the annotations present in the obtained metadata.

#### 2.1.2 Liver hepatocyte-endothelial pair data

The scRNAseq count matrix of carefully isolated hepatocyte-endothelial cell pairs from Halpern et al. [20] were obtained from the GEO database (GSE108561).

#### 2.1.3 Small intestinal scRNAseq datasets

The singlet and multiplet (composing partially dissociated cell clumps) scRNAseq count matrices and the associated metadata (including cell labels for the singlet data) from Andrews et al. [19] were obtained from the GEO database (GSE175664). In addition, a multiplet scRNAseq count matrix comprising partially dissociated cell clumps from another study [18] was used (Manco; GSE154714).

#### 2.1.4 Lung scRNAseq datasets

The singlet and multiplet scRNAseq count matrices and the associated metadata (including cell labels for the singlet data) from Andrews at al. [19] were obtained from the GEO database (GSE154714).

#### 2.1.5 Ovarian cancer scRNAseq and spatial transcriptomics datasets

The scRNAseq count matrix and the associated metadata, including cell labels, from ovarian cancer study by Denisenko et al. [24] were obtained (GSE211956). This also included the spatial transcriptomics data of 8 ovarian cancer tissue sections containing spot-level count matrix, spatial imaging data and tissue coordinates. For scRNAseq, the data was processed to include only genes that were expressed in a minimum of 3 cells and only cells with at least 500 genes, less than 15% mitochondrial genes, and greater than 500 total RNA counts. After the preprocessing step and UMAP visualization, cells were assigned cell type labels by transferring the annotations present in the obtained metadata. For spatial transcriptomics, the individual matrices and imaging data were loaded to create 8 spatial objects. The data were normalized and transformed using the SCT method. Next, PCA was performed on each spatial object following the identification of the top 3000 HVGs. Spots were then clustered by nearest neighbor clustering approach, followed by projection and visualization on UMAP plots of the first 30 PCs. Spot-level deconvolution was performed using the robust cell type decomposition (RCTD) method from the Spacerx R package [25] with the annotated scRNAseq data as reference set.

#### 2.1.6 Breast cancer scRNAseq and spatial transcriptomics datasets

The scRNAseq count matrix and the associated metadata of a breast cancer study, including cell labels, from Wu et al. [26] were obtained from the GEO database (GSE176078). This also included the spatial transcriptomics data of 6 breast cancer tissue sections containing spot-level count matrix, spatial imaging data and tissue coordinates. For scRNAseq, the data were preprocessed, and cells were assigned cell type labels by transferring the annotations present in the obtained metadata. For spatial transcriptomics, the individual matrices and imaging data were loaded to create 6 spatial objects. The data were normalized and transformed using the SCT method. Next, PCA was performed on each spatial object following the identification of the top 3000 HVGs. Spots were then clustered by nearest neighbor clustering approach, followed by projection and visualization on UMAP plots of the first 30 PCs. Spot-level deconvolution was performed using the robust cell type decomposition (RCTD) method from the Spacerx R package [25] with the annotated scRNAseq data as reference set.

### 2.2 The univariate linear model (ULM) approach

#### 2.2.1 Gene signature generation

Cell type specific gene signatures for the full scRNA-seq datasets were generated for each annotated cell type using the FindAllMarkers function of the Seurat R package (version 5.1.0) with a one-versus-all differential gene expression analysis using the default parameters (Wilcoxon rank sum test, minimum log2 fold-change threshold = 0.25, minimum detection fraction = 10%). For each cell type, the differentially expressed genes were ranked based on their fold changes and the top 100 genes with adjusted p values < 0.05 were selected as gene signatures.

#### 2.2.2 Gene signature scoring

The Univariate Linear Model (ULM) was used to calculate gene signature activity scores by modeling the relationship between gene expression profiles for each cell and the cell type signature weights. This analysis was performed using the run_ulm function from the decoupleR R package [27]. For each cell, a t-value was derived from the fitted model, representing the activity score of the respective gene signature. For downstream analysis, only positive activity scores greater than 1 and p-values < 0.05 were considered. Cells with active enrichment scores were classified into two categories: (i) singlets: cells showing only one cell type specific gene signature, indicating a single-cell identity. (ii) doublets and multiplets: cells exhibiting active enrichment for more than one cell type gene signature, representing potential doublets or multiplets.

#### 2.2.4 Multiplet filtering and network plot

Following cell barcode classification, multiplets were grouped by cell type compositions, and the frequency of each multiplet type was computed. To reduce noise, only multiplet types with frequency greater than 10 were considered. The multiplet types were decomposed by cell pairs and frequencies into nodes and edges of a network using the igraph R package [28]. Cell-cell interaction networks were plotted from the nodes and edges using the ggraph R package [29], depicting a physical interaction network inferred from the scRNAseq dataset. The network edges are undirected and are weighted by the frequencies of the cell type pairs (node-node pairs).

### 2.3 Cell pairs predictions and benchmarking

The ULM performance was tested against two other similar methods, Neighborseq [3] and CIcADA [30], using FACS sorted doublet datasets of DC-T and endothelial-hepatocyte pairs with known constituents as ground truths. For the DC-T doublet pairs, DC-specific and T cell specific gene signatures were derived by performing a DC vs T cell DEG on the singlet DC and T cell datasets. Then, ULM prediction was performed on the merged dataset of DC and T cell singlets and DC-T doublets as explained earlier. For Neighborseq prediction, a training gene expression matrix was prepared from the singlet DC and T cell data, using the prep_cell_mat function of the Neighborseq R package with the default parameters (logfc.threshold =1, max.cells = 200, min.pct = 0.25, topn = 50, res = 0.8). Artificial multiplets were generated from the prepared singlet data using the artificial_multiplet function, setting n = 100, to simulate 100 artificial doublets. A random forest model was then trained on the singlets and artificial multiplets using the multiplet_rf function. The trained random forest model was finally used to predict a merged dataset of singlets and DC-T doublets using the pred function which assigned labels to each cell as singlets (DC or T cell) or doublets (DC-T). For CIcADA, the CAMML R package was used. Firstly, gene signatures were built from the singlet DC and T cells using the BuildGeneSets with the default parameters (cutoff.type = ‘logfc’, cutoff = 2, species = ‘Mm’, weight.type = ‘logfc’). Next, predicted gene signature scores were assigned to each cell in the merged dataset of unlabeled singlets and doublets using the CAMML function. This assigned signature scores between 0 and 1 to each cell and cell classifications were performed using a threshold of 0.5. Cells scoring a minimum of 0.5 for two signatures were classified as doublets while those scoring a minimum of 0.5 for only one signature were labeled singlets. The predictive accuracies of the 3 methods (ULM, Neighborseq and CIcADA) were compared using the ConfusionMatrix function of the Caret R package.

For the hepatocyte-endothelial doublets, mouse hepatocyte and endothelial cell signatures were generated by obtaining the mouse hepatocytes and endothelial markers from the PangloDB database (https://panglaodb.se/) since the hepatocyte-endothelial dataset did not have an accompanying annotated singlet dataset. These signatures were then used to predict each cell in the hepatocyte-endothelial doublet dataset using the ULM pipeline as discussed earlier. For Neighborseq predictions, a training gene expression matrix was generated from a hepatocyte and endothelial singlet dataset subsetted from a mouse liver scRNAseq data generated from another study [31]. The hepatocyte-endothelial doublets were then predicted using the same parameters and following the same steps as the DC-T doublets described above. Similarly, for CIcADA, gene signatures were built from all cells in the annotated mouse liver scRNAseq data [31] using the BuildGeneSets function of the CAMML package and the default parameters as stated above. The hepatocyte and endothelial gene signatures were subsetted for downstream analysis. Then, gene signature scoring and cell type assignments were performed for the hepatocyte-endothelial doublets using the CAMML function and a threshold of 0.5 respectively.

### 2.6 Ligand receptor analysis

To assess whether predicted doublets co-expressed ligand-receptor gene pairs, predicted doublet pairs of B/plasma cells and T cells, named T-B doublets, obtained from ULM predictions of the breast cancer scRNAseq dataset were tested. A list of ligand-receptor pairs was obtained from Ramilowski et al. [32] containing 708 unique ligands and 691 unique receptors. For each T-B doublet pair, a ligand or receptor gene was considered expressed if its expression level in that doublet was higher than its mean expression across all T-B doublets. Therefore, a ligand-receptor pair (LRP) was assumed to be co-expressed in a doublet if both ligand and receptor genes were expressed in that doublet. The LRPs which were co-expressed in at least 5 doublets were retained. Next, the co-expressed LRPs were validated for possible colocalization in the matched spatial transcriptomics data of the same breast cancer tissue. Firstly, each spatial spot was tested for the colocalized enrichment of B cells/plasmablasts and T cells. Spots which simultaneously contained at least 5 % B/plasma cells and at least 5 % T cells were considered colocalized T-B spots. The T-B spatial spots were then tested for colocalization of the LRPs that were co-expressed in the T-B doublets. To this end, for each spatial T-B spot, a ligand or receptor gene was considered expressed if the expression level was higher than the average expression across all spatial T-B spots. Similarly, an LRP is considered colocalized in a spot if both ligand and receptor genes are expressed in that spot. Finally, the proportion of T-B spots having the colocalization of the LRPs that were expressed in the T-B doublets was computed.

## 3.0 RESULTS

### 3.1 Predicting cell pairs using ULM and benchmarking

Doublets in scRNAseq contain 2 cells and can be identified by the expression of markers for dual cell type lineages. This concept was shown in a scRNAseq study with single-positive DCs and T cells, and doublet pairs of DC-T cells obtained from co-cultured DCs and T cells (in vitro) or from the lymph nodes, representing ongoing antigen presentation (in vivo) [23]. As shown in Figure 1a, the single-positive T cells and DCs form distinct clusters, with DC-T doublet cluster localized between these populations. Further, the doublets express T-cell specific marker genes such as CD3D, CD3E, CD3G and CD4 (Figure 1b) and DC specific markers such as CD80, CD86, CD83 and CLEC7A (Figure 1b). We also showed this dual marker expression in a dataset containing doublet pairs of hepatocyte-endothelial cells, obtained by FACS sorting of physically connected mouse hepatocytes and endothelial cells. As seen in Figure 1c, the hepatocyte-endothelial doublets co-expressed hepatocyte specific marker genes (Ahr, Cps1, Gls2, Ass1, Fcna, Hhex, Gck, Cyp7a1 and Glul) and endothelial cell specific marker genes (Fcgr2b, C1qtnf1, Flt4, Fabp4, Icam1 and Cldn1). The co-expression of these cell type-specific markers in doublets provides an opportunity for identifying and distinguishing multiplets from other cells.

**Figure 1.**
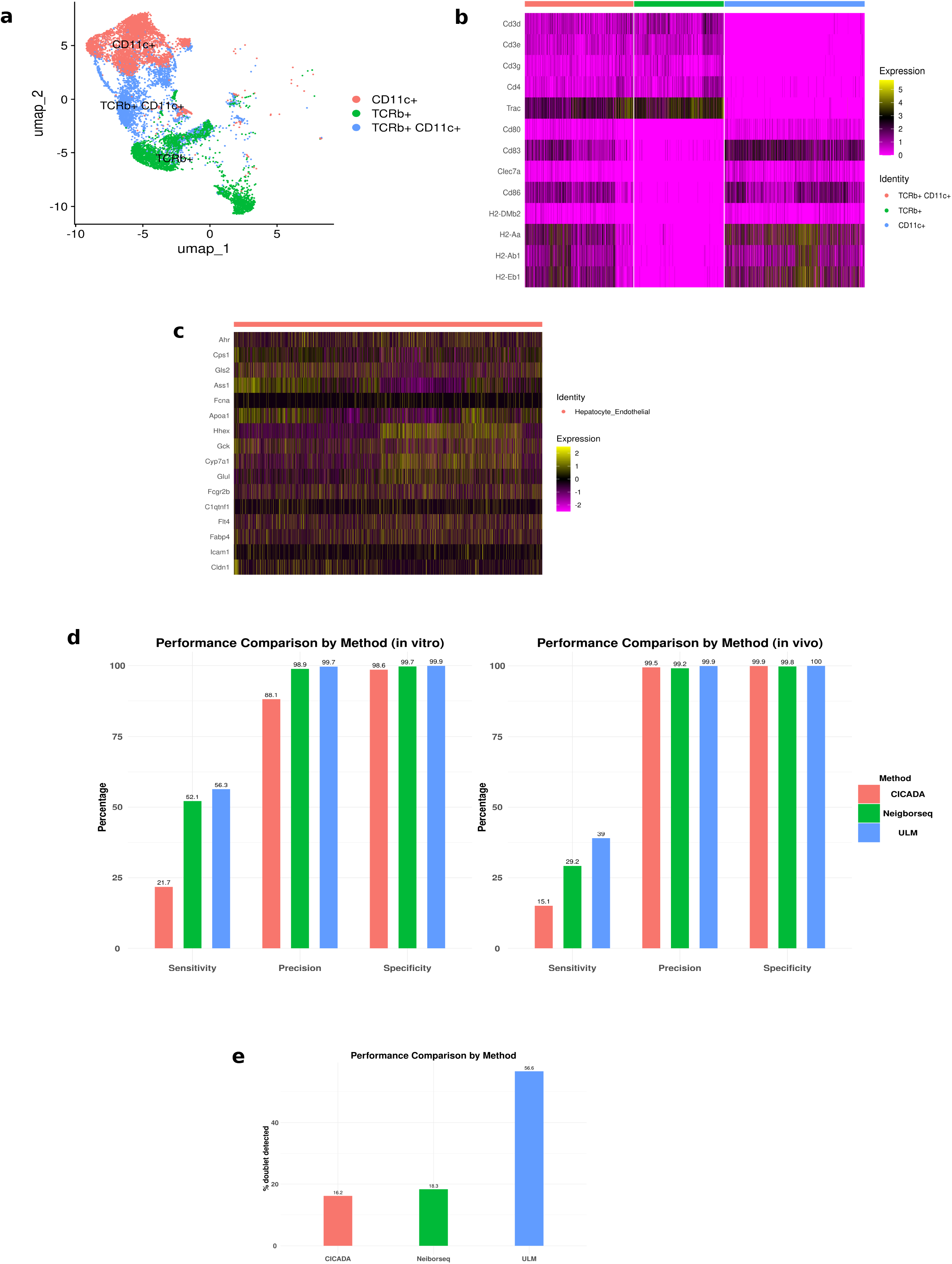
Marker expression, model performance and benchmarking. Using FACS sorted datasets consisting of double-positive dendritic cells and T cells doublets (T-DC) and hepatocyte-endothelial doublets, we showed that doublets express dual lineage-specific cell type markers. The performance of the ULM model was tested on the FACS sorted cell pairs and was compared to two other existing doublet identification/deconvolution models (Neighbor-seq and CICADA). (a) UMAP plot showing distinct clusters formed by dendritic cells (CD11c+), T cells (TCRb+) and T-DC doublets (CD11c+ TCRb+) (b) Heatmap showing the expression of key T cell and dendritic cells markers by T-DC doublets and their singlet counterparts. The normalised gene expression counts were utilised. (c) Heatmap showing the expression of key hepatocyte and endothelial markers by hepatocyte-endothelial doublets. The scaled gene expression values were used. (d) Left: performance metrics for ULM, Neighborseq and CICADA models on a publicly available dataset consisting of FACS sorted single-positive dendritic cells, single-positive T cells and in vitro co-cultured T-DC doublets. Right: Performance metrics for ULM, Neighborseq and CICADA models on a publicly available dataset consisting of sorted dendritic cells, T cells and T-DC doublets isolated from the lymph nodes and spleen of mice (in vivo derived cells) (e) Performance metrics for ULM, Neighborseq and CICADA models on a publicly available dataset consisting of sorted hepatocyte-endothelial cell doublet from partially digested liver tissue.

To distinguish singlets from multiplets and deconvolute the cellular composition of the identified multiplet in single cell RNA-seq data, we propose a new gene signature-based computational framework utilizing a Univariate Linear Model (ULM).

The steps involved in the ULM approach are summarized below:

1. Generating cell type specific signatures from annotated scRNAseq input data
2. Fitting a ULM over the gene expression of each barcode in the data
3. Scoring each cell for cell type-specific signatures and filtering positive enrichment (activity) scores
4. Classifying each barcode based on signature enrichment and selecting multiplets
5. Plotting physical cell-cell interaction networks from predicted multiplet data

We applied the ULM approach to predict doublet cell type composition using two datasets containing FACS-sorted cell pairs and compared its accuracy with two other published methods: Neighborseq [3] and CIcADA [33]. These methods were developed to identify potential doublets and their cell type compositions from scRNAseq datasets. Neighborseq utilizes a random forest model trained on annotated singlets and synthetically generated doublets. On the other hand, CIcADA uses a multi label cell typing approach based on variance-adjusted Mahalanobis scoring to assign cell type probabilities (0-1 scale) for predefined cell type specific gene sets, classifying cells with dual scores above a defined threshold as doublets.

Firstly, we evaluated the predictive accuracy of ULM on the scRNAseq datasets of *in vitro* (co-cultured) and *in vivo* (lymph node derived) dendritic cells-T cells (DC-T) doublets alongside with the singlet DCs and T cells from Gliadi et al. [23]. The *in vitro* dataset comprises 4130 DCs, 2641 T cells and 3203 DC-T doublets, while the *in vivo* dataset contained 1985 DCs, 2081 T cells and 3674 DC-T doublets. The aim here was to test whether the model could correctly predict and identify doublets in a mixed dataset of doublets and singlets. Since the cells were sorted by surface markers prior to single-cell sequencing, the cell type labels and the number of cells (singlet or doublet) were known *a priori* and serve as ground truths. We used the singlet gene expression data to build cell type specific gene signatures which were then used to predict the cell type composition of each cell in a merged dataset of both singlets and doublets using ULM. Barcodes exhibiting enrichment for either the DC or T cell gene signature were classified as predicted singlets, whereas those with positive activity scores for both DC and T cell signatures were identified as doublets. For *in vitro*-derived cells, the model achieved a sensitivity of 56%, implying that it successfully identified and classified 56% of the doublets as DC-T doublets (Figure 1d, Supplementary Table 1). Additionally, the model displayed a remarkable precision of 99.7%, indicating that cells classified as doublets are almost exclusively true doublets, with minimal misclassification of singlets. The specificity was 99.9%, which implied that almost all singlets in the data were identified and classified as singlets (Figure 1d). Given that this method was specifically developed for multiplet detection, sensitivity and precision are the more critical metrics. Interestingly, the model outperformed existing published methods, including Neighborseq [3] and CIcADA [33]. Neighborseq achieved a sensitivity of 52.1%, with slightly lower precision of 98.9% and specificity of 99.7% (Figure 1d). CIcADA performed significantly worse, achieving a sensitivity of only 21.7% with a precision of 88.1% and specificity of 98.6% (Figure 1d). These results underscore the enhanced performance of the proposed ULM method in detecting and classifying doublets effectively.

For the in vivo derived cell populations, ULM achieved a 39% sensitivity with a precision of 99.9% and specificity of 100% (Figure 1d). The remarkably high precision indicates that the model reliably classified true doublets without misclassifying singlets as doublets. The reduced sensitivity likely reflected the increased complexity of the *in vivo* system compared to the *in vitro* system. Despite this limitation, ULM outperformed both Neighborseq and CIcADA. Neighborseq achieved only 29.2% sensitivity with a comparable precision of 99.2% and specificity of 99.8%, while CIcADA identified just 15.1% of doublets with a comparable precision of 99.5% and specificity of 99.9% (Figure 1d). Although all the 3 tools showed remarkably high precision and specificity, ULM showed a superior sensitivity. These findings highlight ULM’s enhanced capability for doublet detection in more complex biological systems.

Secondly, we evaluated the ULM approach on the dataset of hepatocyte-endothelial doublet pairs consisting of 4621 FACS sorted double-positive cells for hepatocyte and endothelial markers from partially dissociated liver tissues [20]. As this dataset lacked an annotated singlet signature, we generated mouse hepatocyte and endothelial cell signatures by obtaining the mouse hepatocytes and endothelial markers from the PangloDB database (https://panglaodb.se/), yielding 138 hepatocyte and 189 endothelial marker genes. These gene signatures were used to predict the possible cell type compositions of the paired hepatocyte-endothelial doublet cell barcodes. Barcodes displaying enrichment exclusively for either hepatocyte or endothelial gene signatures were classified as singlets, whereas those exhibiting positive activity scores for both signatures were designated as doublets. We computed the proportion of barcodes that were predicted as hepatocyte-endothelial doublets and benchmarked this against Neighborseq and CIcADA. Our method successfully identified 56.6% of the hepatocyte-endothelial doublets, substantially outperforming both Neighborseq and CIcADA, which detected only 18.3% and 16.3% of the doublets, respectively (Figure 1e, supp table 1).

Taken together, these results indicate that the ULM approach can successfully identify doublets from scRNAseq datasets with minimal misclassification of singlets as doublets. The method consistently achieves higher sensitivity and precision, outperforming or matching the performance of other published methods.

### 3.2 Predicting multiplets and their cell compositions from partially dissociated tissues

We applied the ULM approach to predict the cellular composition of real multiplets derived from partially dissociated tissues which were subsequently sequenced in studies aimed at deciphering the physical interaction landscape of tissue architectures. Specifically, we analyzed intestinal tissue clumps from Andrews et al. [19], comprising 3671 cells, Manco et al. [18] with 6815 cells, as well as lung tissue clumps from Andrews et al. [19] with 4729 cells.

Using the intestinal multiplet datasets, we first generated intestinal cell type specific gene signatures from the annotated conventional scRNAseq singlet data (Figure 2a) obtained from Andrews et al. [19]. These signatures were then used to predict cell-cell interactions in the two intestinal multiplet datasets using the ULM method. We differentiated between predicted singlets and multiplets based on their enrichment scores: barcodes with a score for one gene signature were classified as singlets, while those with scores for multiple signatures were designated as multiplets, indicating the co-presence of multiple cell types within individual droplets. Application of the ULM algorithm to the Andrews et al. intestinal multiplet dataset revealed approximately 1000 cells as multiplets, comprising 904 doublets, 93 triplets, and 2 quadruplets. Analysis of the Manco et al. [18] intestinal multiplet data yielded substantially more multiplets, with 5,082 cells successfully predicted as containing multiple cell types (1,755 doublets, 2,735 triplets, 522 quadruplets, and 67 pentuplets).

**Figure 2.**
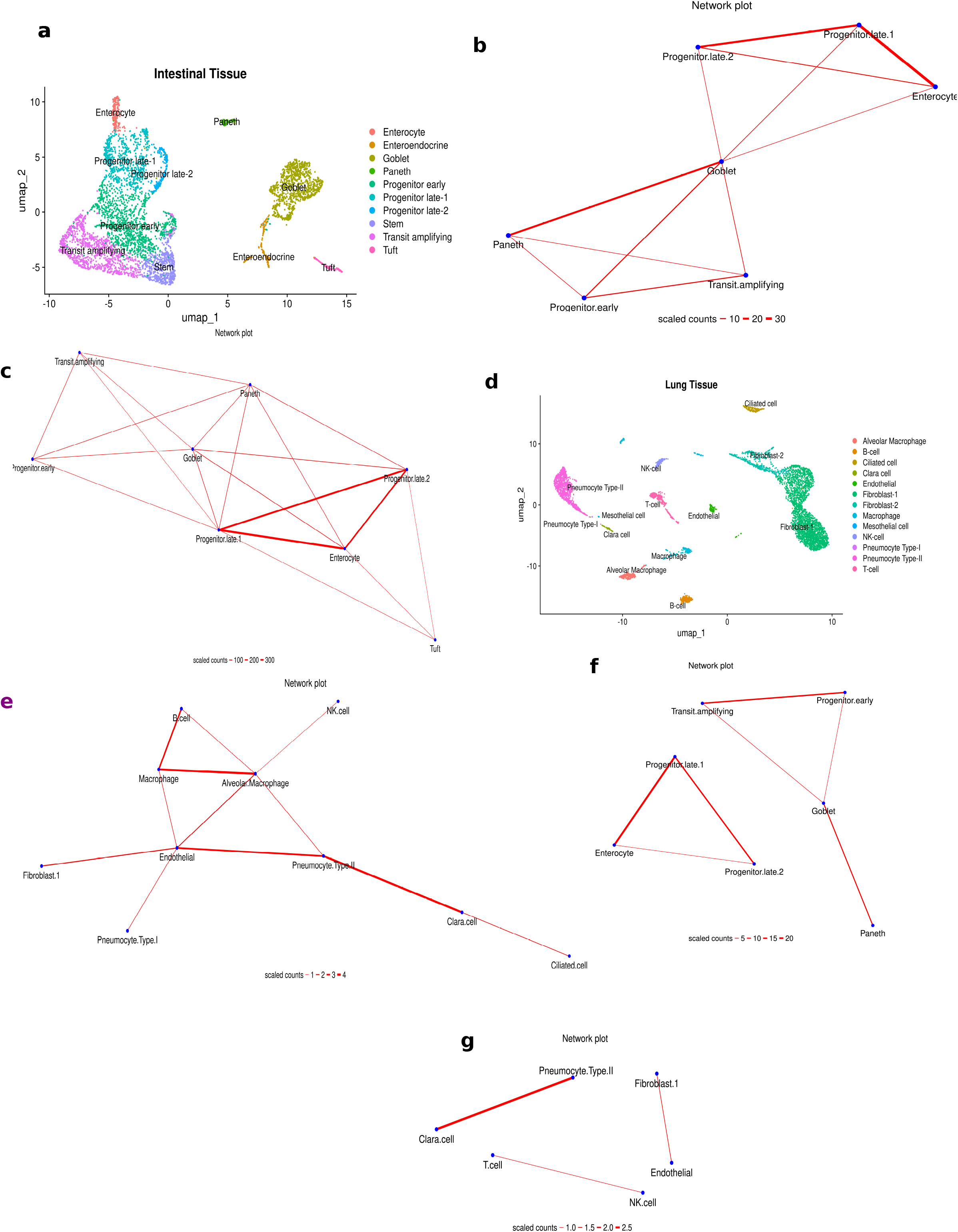
Multiplet predictions from partially digested tissues and conventional scRNAseq “singlet” data. Publicly available datasets of intestinal and lungs multiplets obtained by partial dissociation were utilised. The intestinal and lung singlet scRNA seq data were used to generate signatures to be applied to identify and deconvolute the respective multiplets. (a) UMAP plots of the intestinal singlet scRNAseq data showing cell clustering by annotated cell types. (b-c) network plots of identified multiplets from two datasets of partially digested intestinal tissues. Using the signature generated from the cell types in Figure 3a, the model was applied to two partially dissociated intestinal multiplet datasets without prior annotation. The identified multiplet types were decomposed to generate a network plot of physical cell interactions. Interestingly, known microanatomical structure of the small intestine. Each multiplet type has significantly positive enrichment scores (threshold: p < 0.05 & score >1) for two or more cell signatures and has a count of at least 10. The edge counts were scaled by 10. (d) UMAP plots of the lung singlet scRNAseq data showing cell clustering by annotated cell types. (e) Network plot of multiplets identified from a dataset of partially digested lung tissues. Using the signature generated from the cell types in Figure 3d, the model was applied to a partially dissociated lung multiplet datasets without prior annotation. The identified multiplet types were decomposed to generate a network plot of physical cell interactions. The microanatomical structure depicting the stromal and functional units of the lung epithelium were recapitulated by the network. Also, there was a small network of immune cell interactions. Each multiplet type has significantly positive enrichment scores (threshold: p < 0.05 & score >1) for two or more cell signatures and has a count of at least 10. The network edge counts were scaled by 10. (f) Network plots of predicted significant multiplets from standard scRNAseq intestinal datasets (in 3a). Interestingly, all the interactions identified were present in the multiplet network (in 3e), indicating the biological plausibility of the identified multiplet types. The edge counts were scaled by 10. (g) Network plots of predicted significant multiplets from standard scRNAseq lung datasets (in 3d). Interestingly, two of the three interactions identified were present in the multiplet network (in 3e), indicating the biological plausibility of the identified multiplet types. The edge counts were scaled by 10.

To visualize cell-to-cell interactions, we constructed physical interaction networks from the predicted multiplets, focusing on multiplet types occurring in more than seven cells. The resulting networks (Figures 2b-c) recapitulated the known microanatomical structure of the small intestine [34,35]. The networks were dominated by interactions involving intestinal progenitors and enterocytes, consistent with the dense population of absorptive enterocytes in the intestinal epithelia that are constantly renewed by newly differentiated cells from progenitors. Furthermore, there were interactions of the transit amplifying cells, early progenitors and Paneth cells, also indicating a proliferative niche with regenerative potential to the fully differentiated Paneth cells. Goblet cells were centrally placed, having physical connections with most other cells in the network. This recapitulates the known distribution of the mucus-secreting goblet cells along the entire length of the intestine, where they are strategically located among other cells and are also formed from proliferative cells (progenitors and transit amplifying cells). Additionally, Figure 2c (generated from Manco et al. data) displayed a more complex network, also featuring the immunological tuft cells interacting with enterocytes and the progenitors. This can be explained by their distribution along the intestine where they are interspersed with enterocytes, as well as their differentiation from progenitors [36].

For the lung multiplet data prediction comprising 4729 cells, we generated cell type specific gene signatures from the annotated conventional scRNAseq lung singlet data obtained from the same study (Figure 2d). We then applied the ULM pipeline to the multiplet dataset to predict their cellular composition and plot a physical interaction network (Figure 2e). Again, we differentiated between predicted singlets and multiplets based on their enrichment scores: barcodes with a score for one gene signature were classified as singlets, while those with scores for multiple signatures were classified as multiplets. In contrast to the intestinal datasets, we only identified 377 multiplets in the lung tissue, of which 366 were predicted as doublets and only one as triplet. This marked reduction in multiplet frequency may be attributed to the less continuous organization of lung tissue compared to the intestinal epithelium, which features more tight junctions that could facilitate the formation of multicellular aggregates during tissue dissociation. A large proportion of the cells in the lungs are migratory/resident immune cells as well as vascular endothelial cells which are less tightly and non-continuously connected with epithelial cells. Nonetheless, physical interaction networks were plotted from the predicted multiplets using only multiplet types occurring in more than 10 barcodes. The predicted cellular network effectively captured the known structural and functional organization of the lung epithelium (Figure 2e). The cell-cell interactions involving endothelial cells, pneumocytes and alveolar macrophages recapitulate the alveolar unit of the lungs, representing the functional unit of the lungs involving gaseous exchange [37]. Also, an interaction between ciliated and Clara cells partially captures the small and terminal bronchiolar unit of the lungs [38]. Finally, there was a localized immune network that features interactions between immune cells (Figure 2e). Overall, these results emphasize the applicability of the ULM approach to predict cellular interaction networks from tissues.

### 3.4 Identifying and predicting multiplets in conventional scRNAseq

Finally, we applied the ULM approach to conventional scRNAseq datasets as it was specifically being developed for this task. We tested 4 scRNAseq datasets for this purpose. We initially tested the model on datasets of healthy lung and intestinal tissues from Andrews et al. [19], and then on 2 cancer scRNAseq dataset from ovarian cancer [24] and breast cancer [26] tissues. For each scRNAseq dataset, we applied the complete ULM pipeline, which briefly involved gene signature generation from cell annotation, ULM scoring per barcode, cell type classification and network plots from predicted multiplets. For the intestinal scRNAseq data comprising 5279 cells, we used the annotated clusters (Figure 2a) to generate cell signatures which were then used to score and classify each cell barcode in the dataset (supp table 2 & 3). We identified 698 multiplets, of which 673 were predicted as doublets and 25 were predicted as triplets. The predicted multiplets were used to plot a physical interaction network (Figure 2f). Interestingly, we uncovered multiple cell-cell interactions which mimicked the network obtained from the partially dissociated intestinal multiplets (Figures 2b-c), more like a subnetwork. For example, the progenitor-enterocyte interactions and the goblet-progenitor-transit amplifying interactions which dominated the networks in Figures 2b & 3c (partially dissociated) were enriched in Figure 2f (conventional scRNAseq). Similarly, for the lung scRNAseq data comprising 6084 cells, we generated cell type specific gene signatures from the annotated clusters (Figure 2d). These were then used to score and classify each cell barcode in the scRNAseq data (supp table 2 & 3). We identified 120 multiplets, all of which were predicted as doublets. We plotted a cell-cell interaction network from the predicted multiplets. We uncovered some isolated cell interactions (Figure 2g) which partially mimicked the interactions found in the partially dissociated lung multiplet network (Figure 2f). For example, endothelial-fibroblast and Clara-ciliated cell interactions were obtained from the scRNAseq data which were present in the partially dissociated lung multiplet network (Figure 2e). These results implied that biologically meaningful interactions could be generated from conventional scRNAseq data of healthy tissue. Indeed, these represent the potentially undissociated cell fractions since the connections were also found in actual multiplet datasets.

Next, for the breast cancer scRNAseq data comprising ∼100,000 cells, we generated gene signatures from the annotated clusters shown in Figure 3a. We employed the signatures in the model to score and assign predicted cell classes to each barcode in the data (Supp Tables 2 & 3). We identified 1624 multiplets, of which 1621 were predicted as doublets and 3 were predicted as triplets. The predicted multiplets were filtered and utilized to generate a physical interaction network plot (Figure 3b). The resulting network (Figure 3b) featured multiple immune cell interactions as well as a stromal unit involving stromal cell interactions such as cancer associated fibroblasts (CAFs), perivascular-like cells (PVL) and endothelial cells. Cancer cells were less involved in physical interactions as they cluster away from other cells, being connected only with plasmablasts. We validated this result in the matched ovarian cancer spatial transcriptomics data from the same study consisting of ∼16,000 Visium spatial spots from six breast cancer tissue sections. We deconvoluted the cellular composition of each spatial spot using the RCTD method with the matched scRNAseq data as reference. Cell-cell colocalization was then assessed by Spearman’s correlation, such that cells which colocalize in spatial spots would possess positive correlation coefficients (r). Logically, cells which colocalize in spatial spots are in close proximity in tissues and hence have a high likelihood of being involved in physical interactions. Interestingly, the enriched cell-cell interactions obtained from the scRNAseq data (Figure 3b) showed positive correlations in the spatial spots (Figure 3c). For example B cell-T cell (r = 0.5), B cell-plasmablasts (r = 0.15), CAF-endothelial (r = 0.24), CAF-myeloid (r = 0.39), CAF-PVL (r = 0.17), endothelial-PVL (r = 0.5), endothelial-T cell (r = 0.24), and myeloid-T cell (r = 0.45) interactions showed statistically significant positive correlations in spatial spots and were concurrently enriched in the ULM network. Also, cancer cells showed very strong negative correlation with all other cells, showing the least negative (highest) correlation with plasmablasts. This explains why the cancer cells cluster away from other cells in the ULM network with connection only with plasmablasts. This reinforces the accuracy (sensitivity) of the ULM approach.

**Figure 3.**
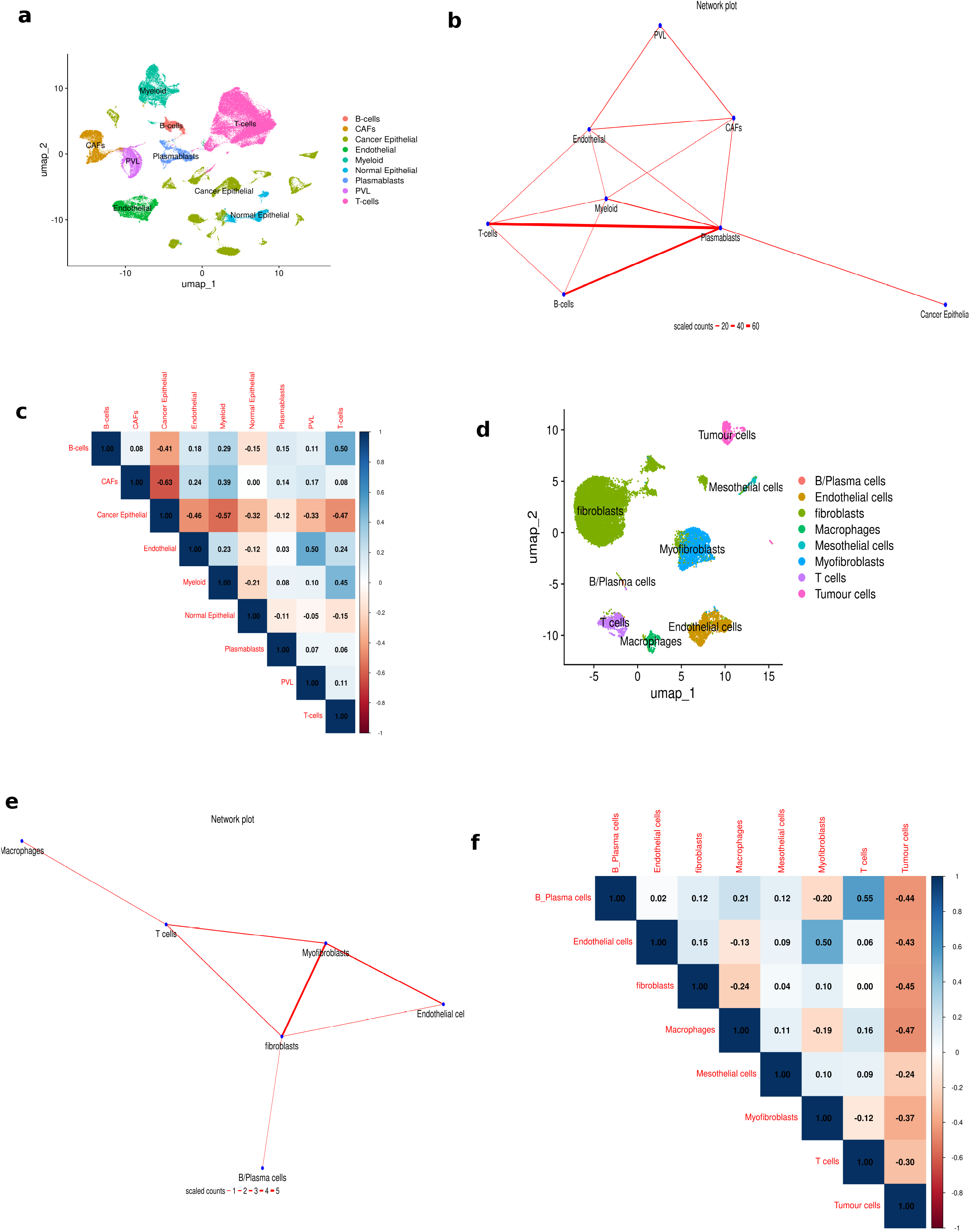
predictions of scRNAseq cancer datasets and validation by spatial transcriptomics. (a) UMAP plots of the breast cancer scRNAseq data showing cell clustering by annotated cell types. (b) Network plot of significant multiplet types inferred from the breast cancer scRNAseq data (in 4a). Important immune and stromal network clusters were depicted. Each multiplet type had significantly positive enrichment scores (threshold: p < 0.05 & score >1) for two or more cell signatures and a count of at least 10. The edge counts were scaled by 10. (c) Spatial spot level correlation coefficients of cell types inferring cell-cell spatial colocalization. The associated spatial transcriptomics data obtained from the same breast cancer tissue as the scRNAseq in 4a was processed and spot level deconvolution was performed to identify cell types present in each spatial spot. Correlation analysis between spot level cell type scores was used to identify spatially co-localizing cell types. As shown, most of the interaction edges identified in 4b had positive correlation coefficients. Also, cancer cells negatively correlated with all other cells and this reasonably explains why they clustered away from other cells in the network plot. All correlation coefficients are statistically significant (p < 0.05). d) UMAP plots of ovarian cancer scRNAseq data showing cell clustering by annotated cell types. (e) Network plot of significant multiplet types inferred from the ovarian cancer scRNAseq data (in 4d). Immune and stromal interactions were enriched in the network. Each multiplet type has significantly positive enrichment scores (threshold: p < 0.05 & score >1) for two or more cell signatures and has a count of at least 5. The edge counts were scaled by 10. (f) Spatial spot level correlation coefficients of cell types inferring cell type spatial colocalization. The matched spatial transcriptomics data obtained from the same breast cancer tissue as the scRNAseq in 4d was processed and spot level deconvolution was performed to identify cell types present in each spatial spot. Correlation analysis between spot level cell type scores was used to identify spatially co-occurring cell types (spatial colocalization). As shown, most of the interaction edges identified in 4e had positive correlation coefficients. All correlation coefficients are statistically significant (p < 0.05).

Finally, for the ovarian cancer scRNAseq data comprising ∼12000 cells, cell type specific gene signatures were generated from the obtained cluster annotations shown in Figure 3d. We utilized the obtained signatures to score and predict the cellular composition of each barcode in the dataset using the ULM pipeline (supp table 2 & 3). We identified 145 multiplets, all of which were predicted as doublets. The predicted multiplets were filtered and utilized to generate a physical interaction network plot (Figure 3e). The resulting network (Figure 3e) showed enriched immune-stromal interactions such as myofibroblast-fibroblast, macrophage-T cell, fibroblast-endothelial cell, fibroblast-T cell and fibroblast-B cell interactions. Similarly, we validated this result in the matched ovarian cancer spatial transcriptomics data from the same study consisting of ∼21000 Visium spatial spots from nine ovarian cancer tissue sections. The cellular composition of each spatial spot was deconvoluted using RCTD with the matched scRNAseq data as reference, and we assessed cell-cell colocalization by Spearman’s correlation. Interestingly, the enriched cell-cell interactions obtained from the scRNAseq data (Figure 3e) showed positive correlations in the spatial spots (Figure 3f). For example, fibroblast-B cells (r = 0.12), fibroblast-endothelial cell (r = 0.15), myofibroblast-endothelial (r = 0.5), fibroblast-myofibroblast (r = 0.1), and macrophage-T cell (r = 0.16), interactions showed statistically significant positive correlations in spatial spots and were concurrently enriched in the ULM network. Also, tumor cells showed very strong negative correlation with all other cells, hence this explains why they were not enriched in the ULM network, further reassuring the precision of the ULM method.

### 3.5 Downstream analysis: ligand-receptor interaction

One additional way to further validate the accuracy of the ULM predictions is to assess multilineage marker expression and ligand-receptor co-expression in predicted multiplets. We assessed these in doublets formed between T cells and B cell lineage (B cells or plasmablasts) obtained from the breast cancer predicted network. In this regard, we selected doublets of the type B cell-T cell and plasmablast-T cell, representing T-B cell interactions (T-B doublets). We obtained a total of 621 T-B doublets. Interestingly, these cells showed simultaneous expression of both T cell markers such as CD3D, CD3E, CD3G, CD4 and CD8A, and B cell lineage markers such as CD79A, CD79B, JCHAIN, IGKC, CD24 and CD27 (Figure 4a). Also, 130 ligand-receptor pairs (LRP) were co-expressed in at least 5 T-B doublets (supp table 4). It is worth mentioning that a ligand or receptor gene is considered expressed in a given T-B doublet if the expression level of that gene is greater than its average expression across all T-B doublets. Interestingly, the top 50 enriched LRPs featured many immune-related pairs involving human leukocyte antigen (HLA) and T-cell receptor (TCR) interactions such as HLA-C_CD3G, HLA-C_CD3G, HLA-B_CD3D, HLA-B_CD3G, HLA-A_CD3D, HLA-A_CD3G, B2M_CD3D and B2M_CD3G (Figure 4b) which are pointers to potentially ongoing antigen presentation [39,40]. Also, there was a high enrichment of VIM_CD44, expressed in more than 30% of the T-B doublets. VIM is a cytoskeletal protein which is required for efficient antigen presentation in B cells [41], while CD44 has been reported to be upregulated in activated T cells following antigen presentation [42]. Finally, we tested the 130 co-expressed LRPs in the T-B doublets for concurrent enrichment in spatial spots where T cells and B cells are colocalized (T-B spots). We first identified a total of 1173 colocalized T-B spots as spatial spots containing at least 5% T cells and 5% either of B cells or plasmablasts. We then assessed the 130 enriched LRPs from the T-B doublets for colocalization (co-expression) in the T-B spots. Again, an LRP of interest was considered expressed in a T-B spot if both ligand and receptor genes are expressed at a level higher than the average expression across all T-B spots (see methods). Interestingly, 120 of the 130 LRPs were found to be co-localized in more than 10 T-B spots (supp table 4). Also, the top 50 colocalized LRPs in the T-B spots featured HLA-TCR interactions such as HLA-A_CD3D, HLA-B_CD3D, HLA-C_CD3D, and B2M_CD3D, as well as VIM-CD44 interaction (Figure 4c). Taken together, these results suggest that the predicted doublets truly contain 2 cells which are undergoing biologically relevant interactions.

**Figure 4.**
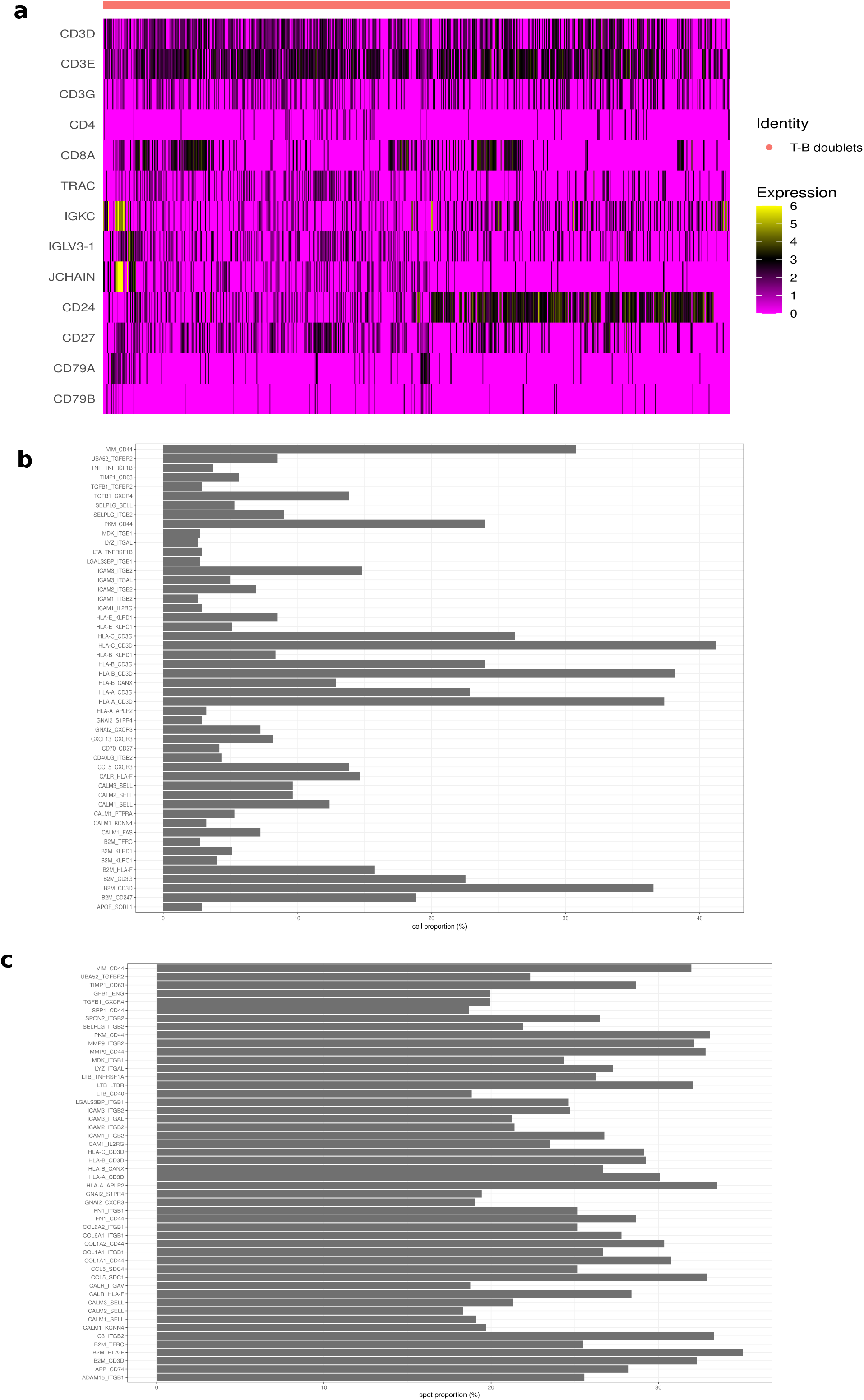
Ligand-receptor analysis. a) Dual lineage marker expression: predicted T-B doublets depicting interactions between T and B cells showed the simultaneous expression of both T cell and B cell markers. b) Top 50 enriched coexpressed LRPs in T-B doublets. Each ligand or receptor gene is considered expressed in a doublet if the expression level in that doublet is greater than the average expression across all doublets. A total of 130 LRPs were coexpressed in at least 5 doublets. The top 50 most expressed LRPs in doublets were displayed. c) Top 50 spatially co-localised LRPs in T-B spatial spots. Visum spots with at least 5% T cells and 5% B cell populations were considered T-B spots. The 130 LRPs that were enriched in doublets were evaluated for colocalization in spatial T-B spots. Each ligand or receptor gene is considered expressed in a T-B spot if the expression level in that spot is greater than the average expression across all T-B spots. 120 of the 130 LRPs showed reasonable colocalization. The top 50 most colocalised LRPs were displayed.

## 4.0 DISCUSSION/CONCLUSION

We developed a new method which utilizes a ULM to detect multiplets (mostly doublets) from scRNAseq data, predict the cellular composition, and infer a physical interaction network from the multiplet distribution. This method leverages the undissociated cell fraction occurring naturally in scRNAseq to make meaningful biological interpretations regarding the ongoing cell interactions in healthy and diseased tissues. Indeed, our method showed good sensitivity with excellent precision in predicting FACS sorted cell pairs with known constituents, recording comparable and/or superior performance over two other existing methods [3,33]. Also, applying it to scRNAseq data of partially dissociated tissues containing real multiplets unraveled physical networks which recapitulated the microanatomical structures of the tissues. This further underscores the accuracy in our predictions to capture biologically meaningful interactions. Finally, we tested our method on classical scRNAseq datasets and obtained biologically reasonable results. When tested on classical cancer scRNAseq datasets, we recovered important interactions which followed biologically plausible cell interactions, validated by cell-cell colocalization in matched spatial transcriptomics datasets. This reassured the accuracy of our method in depicting physical interactions only between cells that were truly in close proximity in tissues. Finally, we assessed captured T-B interactions and found these doublets to show dual lineage marker expression, indicating that they were actual doublets formed between T and B cells rather than falsely predicted interactors. Since the most biologically plausible purpose of this interaction is antigen presentation [39], we showed the enrichment of multiple LRPs involving HLA-TCR interactions which are indicative of potentially ongoing antigen presentation. Interestingly, spatial spots with colocalized T and B cells also displayed a high enrichment of LRPs involved in antigen presentation as well as other LRPs enriched in the T-B doublets. This further reinforces the connection between our predictions from scRNAseq and truly ongoing biological events in tissues.

Our approach has one major limitation. While we successfully applied the ULM approach to identify and deconvolute multiplets (doublets) from scRNAseq data, we could not differentiate random doublets from real (biological) doublets. Random doublets are artificial doublets which result from random combinations of singlets during scRNAseq library preparation where two fully dissociated cells become co-captured in a reaction droplet. These kinds of random doublets are not reflective of tissue interaction landscape and ought to be removed from biological interpretation of tissue dynamics. Real doublets are the true biological doublets that represent an undissociated fraction (cell aggregates) in scRNAseq, which reflect true biological interaction phenomena [22] and are the main targets of the ULM method. Notwithstanding, random doublets can be partially eliminated by assessing known biological plausibility of identified interactions and setting thresholds for acceptable doublet frequencies. In this regard, biologically plausible interactions that are detected at high frequency can be interpreted as potentially real interactions with reasonable levels of confidence. Also, predicted interactions can be validated with an accompanying spatial transcriptomics network constructed from the same tissues to complement ULM predictions and make meaningful interpretations.

## 5.0 DATA AVAILABILITY

All the scRNAseq and spatial transcriptomics datasets utilised in this study were made publicly available by the authors of the cited studies. All scripts and intermediate files associated with this analysis will be made available upon reasonable request through the corresponding author. The ULM package is still being built and will be made publicly available as soon as possible.

## 7.0 ACKNOWLEDGEMENT

This work was supported by Science Foundation Ireland (SFI) through grants 18/SPP/3522 and 22/FFP-A/10729. SAH was supported through the SFI Research Training in Genomics Data Science, grant number 18/CRT/6214.

## SUPPLEMENTARY TABLES

1. Unfiltered scores, final classification and benchmarking statistics of cell pairs
2. Unfiltered enrichment scores for multiplets and singlets scRNAseq barcodes
3. Final classification of multiplets and singlets scRNAseq barcodes

## REFERENCES

1. Boisset J-C, Vivié J, Grün D, Muraro MJ, Lyubimova A, Van Oudenaarden A. Mapping the physical network of cellular interactions. Nat Methods. 2018;15:547–53.

2. Peng L, Wang F, Wang Z, Tan J, Huang L, Tian X, et al. Cell–cell communication inference and analysis in the tumour microenvironments from single-cell transcriptomics: data resources and computational strategies. Brief Bioinform. 2022;23:bbac234.

3. Ghaddar B, De S. Reconstructing physical cell interaction networks from single-cell data using Neighbor-seq. Nucleic Acids Res. 2022;50:e82.

4. Andrews TS, Hemberg M. Identifying cell populations with scRNASeq. Mol Aspects Med. 2018;59:114–22.

5. Dimitrov D, Türei D, Garrido-Rodriguez M, Burmedi PL, Nagai JS, Boys C, et al. Comparison of methods and resources for cell-cell communication inference from single-cell RNA-Seq data. Nat Commun. 2022;13:3224.

6. Efremova M, Vento-Tormo M, Teichmann SA, Vento-Tormo R. CellPhoneDB: inferring cell–cell communication from combined expression of multi-subunit ligand–receptor complexes. Nat Protoc. 2020;15:1484–506.

7. Jin S, Guerrero-Juarez CF, Zhang L, Chang I, Ramos R, Kuan C-H, et al. Inference and analysis of cell-cell communication using CellChat. Nat Commun. 2021;12:1088.

8. Browaeys R, Saelens W, Saeys Y. NicheNet: modeling intercellular communication by linking ligands to target genes. Nat Methods. 2020;17:159–62.

9. Wang Y, Wang R, Zhang S, Song S, Jiang C, Han G, et al. iTALK: an R package to characterize and illustrate intercellular communication. BioRxiv. 2019;507871.

10. Cabello-Aguilar S, Alame M, Kon-Sun-Tack F, Fau C, Lacroix M, Colinge J. SingleCellSignalR: inference of intercellular networks from single-cell transcriptomics. Nucleic Acids Res. 2020;48:e55–e55.

11. Ahmed R, Zaman T, Chowdhury F, Mraiche F, Tariq M, Ahmad IS, et al. Single-cell RNA sequencing with spatial transcriptomics of cancer tissues. Int J Mol Sci. 2022;23:3042.

12. Williams CG, Lee HJ, Asatsuma T, Vento-Tormo R, Haque A. An introduction to spatial transcriptomics for biomedical research. Genome Med. 2022;14:68.

13. Longo SK, Guo MG, Ji AL, Khavari PA. Integrating single-cell and spatial transcriptomics to elucidate intercellular tissue dynamics. Nat Rev Genet. 2021;22:627–44.

14. Tirado-Lee L. A more precise way to find the needle in the haystack: Identifying rare biology with Xenium In Situ [Internet]. [cited 2024 Aug 27]. Available from: https://www.10xgenomics.com/blog/a-more-precise-way-to-find-the-needle-in-the-haystack-identifying-rare-biology-with-xenium-in-situ

15. Kalaimani A, Kim A, Berridge J, Nguyen B, Mohabbat S, Gantt R, et al. 92 Characterization of the tumor microenvironment using the xenium in situ platform and FFPE tissue array technology [Internet]. BMJ Specialist Journals; 2023 [cited 2024 Aug 27]. Available from: https://jitc.bmj.com/content/11/Suppl_1/A105.abstract

16. Patrick M, He S, Ilcisin S, Kroeppler D, Danaher P, Reeves J, et al. Uncover spatial signatures of tumor microenvironment and oncogenic pathways using 6,000-plex single-cell spatial molecular imaging on FFPE skin squamous cell carcinoma. Cancer Res. 2024;84:3644–3644.

17. He S, Bhatt R, Brown C, Brown EA, Buhr DL, Chantranuvatana K, et al. High-plex imaging of RNA and proteins at subcellular resolution in fixed tissue by spatial molecular imaging. Nat Biotechnol. 2022;40:1794–806.

18. Manco R, Averbukh I, Porat Z, Bahar Halpern K, Amit I, Itzkovitz S. Clump sequencing exposes the spatial expression programs of intestinal secretory cells. Nat Commun. 2021;12:3074.

19. Andrews N, Serviss JT, Geyer N, Andersson AB, Dzwonkowska E, Šutevski I, et al. An unsupervised method for physical cell interaction profiling of complex tissues. Nat Methods. 2021;18:912–20.

20. Halpern KB, Shenhav R, Massalha H, Toth B, Egozi A, Massasa EE, et al. Paired-cell sequencing enables spatial gene expression mapping of liver endothelial cells. Nat Biotechnol. 2018;36:962–70.

21. Hameed SA, Kolch W, Brennan D, Zhernovkov V. Physical cell-cell interactions regulate transcriptional programmes that control the responses of high grade serous ovarian cancer patients to therapy. bioRxiv. 2024;2024–04.

22. Wolock SL, Lopez R, Klein AM. Scrublet: Computational Identification of Cell Doublets in Single-Cell Transcriptomic Data. Cell Syst. 2019;8:281-291.e9.

23. Giladi A, Cohen M, Medaglia C, Baran Y, Li B, Zada M, et al. Dissecting cellular crosstalk by sequencing physically interacting cells. Nat Biotechnol. 2020;38:629–37.

24. Denisenko E, de Kock L, Tan A, Beasley AB, Beilin M, Jones ME, et al. Spatial transcriptomics reveals ovarian cancer subclones with distinct tumour microenvironments. bioRxiv. 2022;2022–08.

25. Cable DM, Murray E, Zou LS, Goeva A, Macosko EZ, Chen F, et al. Robust decomposition of cell type mixtures in spatial transcriptomics. Nat Biotechnol. 2022;40:517–26.

26. Wu SZ, Al-Eryani G, Roden DL, Junankar S, Harvey K, Andersson A, et al. A single-cell and spatially resolved atlas of human breast cancers. Nat Genet. 2021;53:1334–47.

27. Badia-i-Mompel P, Vélez Santiago J, Braunger J, Geiss C, Dimitrov D, Müller-Dott S, et al. decoupleR: ensemble of computational methods to infer biological activities from omics data. Bioinforma Adv. 2022;2:vbac016.

28. Csárdi G, Nepusz T, Müller K, Horvát S, Traag V, Zanini F, et al. igraph for R: R interface of the igraph library for graph theory and network analysis [Internet]. Zenodo; 2025 [cited 2025 Mar 4]. Available from: https://zenodo.org/records/14736815

29. Pedersen TL, RStudio. ggraph: An Implementation of Grammar of Graphics for Graphs and Networks [Internet]. 2024 [cited 2025 Mar 4]. Available from: https://cran.r-project.org/web/packages/ggraph/index.html

30. Schiebout C, Lust HE, Huang YH, Frost HR. Cell type-specific Interaction Analysis using Doublets in scRNA-seq (CIcADA). bioRxiv [Internet]. 2023 [cited 2025 Jan 16]; Available from: https://www.ncbi.nlm.nih.gov/pmc/articles/PMC9949061/

31. Su Q, Kim SY, Adewale F, Zhou Y, Aldler C, Ni M, et al. Single-cell RNA transcriptome landscape of hepatocytes and non-parenchymal cells in healthy and NAFLD mouse liver. Iscience [Internet]. 2021 [cited 2025 Mar 4];24. Available from: https://www.cell.com/iscience/fulltext/S2589-0042(21)01201-3?elqTrackId=20a3bbcb605d4288a7f513942cb6a8b0

32. Ramilowski JA, Goldberg T, Harshbarger J, Kloppmann E, Lizio M, Satagopam VP, et al. A draft network of ligand–receptor-mediated multicellular signalling in human. Nat Commun. 2015;6:7866.

33. Schiebout C, Lust H, Huang Y, Frost HR. Cell type-specific interaction analysis using doublets in scRNA-seq. Bioinforma Adv. 2023;3:vbad120.

34. Kong S, Zhang YH, Zhang W. Regulation of Intestinal Epithelial Cells Properties and Functions by Amino Acids. BioMed Res Int. 2018;2018:1–10.

35. Peterson LW, Artis D. Intestinal epithelial cells: regulators of barrier function and immune homeostasis. Nat Rev Immunol. 2014;14:141–53.

36. Feng X, Flüchter P, De Tenorio JC, Schneider C. Tuft cells in the intestine, immunity and beyond. Nat Rev Gastroenterol Hepatol. 2024;21:852–68.

37. Whitsett JA, Alenghat T. Respiratory epithelial cells orchestrate pulmonary innate 4 immunity. Nat Immunol. 2015;16:27–35.

38. Reynolds SD, Malkinson AM. Clara Cell: Progenitor for the Bronchiolar Epithelium. Int J Biochem Cell Biol. 2010;42:1–4.

39. Pishesha N, Harmand TJ, Ploegh HL. A guide to antigen processing and presentation. Nat Rev Immunol. 2022;22:751–64.

40. Shah K, Al-Haidari A, Sun J, Kazi JU. T cell receptor (TCR) signaling in health and disease. Signal Transduct Target Ther. 2021;6:1–26.

41. Tsui C, Maldonado P, Montaner B, Borroto A, Alarcon B, Bruckbauer A, et al. Dynamic reorganisation of intermediate filaments coordinates early B-cell activation. Life Sci Alliance. 2018;1:e201800060.

42. Schumann J, Stanko K, Schliesser U, Appelt C, Sawitzki B. Differences in CD44 Surface Expression Levels and Function Discriminates IL-17 and IFN-γ Producing Helper T Cells. PLoS ONE. 2015;10:e0132479.

